# Optimization of a Multi-Feature AI Ensemble and Voting System for MAPKAPK2 Inhibitor Discovery

**DOI:** 10.1101/2024.11.26.625342

**Authors:** Hayden Chen

## Abstract

The identification of an effective inhibitor is an essential starting point in drug discovery. Unfortunately, many issues arise with conventional high-throughput screening methods. Thus, new strategies are needed to filter through large compound screening libraries to create target-focused, smaller libraries. Effective computational methods in this respect have emerged in the past decade or so; among these methods is machine learning. Herein, we explore an ensemble Deep Learning model trained on MAPKAPK2 bioactivity data. This ensemble ML model consists of ten individual models trained on different features, each optimized for MAPKAPK2 inhibitor identification. Voting systems were established alongside the model. Using these voting systems, the ensemble model achieved an accuracy score of 0.969 and precision score of 0.964 on a testing set, in addition to reporting a false positive rate of 0.014 on an inactive compound set. The reported metrics indicate an effective initial step for novel MAPKAPK2 inhibitor identification and subsequent drug development, with applicability to other kinase targets.

## 2. Background

### 2.1 Importance of Target

Protein kinases are enzymes that regulate cellular pathways by facilitating the transfer of a phosphate group from ATP to its substrates [1]. Mitogen-activated protein kinase-activated protein kinase 2 (MAPKAPK2) is a downstream target of the evolutionarily conserved p38 mitogen-activated protein kinase (MAPK) signaling pathway, which in humans regulates many cellular processes, including apoptosis, cell division, cell motility, and inflammatory responses to oxidative stress. This pathway is activated by stress, proinflammatory cytokines (chemokines), chronic inflammatory conditions, and cancer [2,3]. Within these processes, MAPKAPK2 has been shown to regulate RNA-binding-proteins (RBPs) [3] which in turn regulate the translation of various cancer-linked mRNA transcripts through posttranscriptional modulation of transcript stability [4]. In the p38 pathway, MAPKAPK2 phosphorylates and inhibits Tristetraprolin (TTP), an RBP that promotes decay of the mRNA transcripts of IL-1β (Interleukin-1β) and TNF-α (Tumor Necrosis Factor-α) [5], which are inflammatory cytokines that play critical roles in tumorigenesis [6,7]. It has been conjectured that MAPKAPK2 activation can contribute to radiation resistance in head and neck cancers [8], implying the value of MAPKAPK2 inhibition in disrupting tumor growth. Moreover, MAPKAPK2 inhibitors (TBX-1 and TBX-2) have been shown to improve metabolism and atherosclerotic plaque stability [9], in addition to lowering blood glucose and improving insulin sensitivity in obesity-induced type 2 diabetes [10]. In the case of rheumatoid arthritis, p38 MAPK inhibitors have been thus far limited by tachyphylaxis, making the downstream target MAPKAPK2 a more optimal target in certain autoimmune diseases [11]. These findings indicate that MAPKAPK2 is a promising therapeutic target in treating various diseases.

### 2.2 AI in Drug Design

Small-molecule kinase inhibitors have been used to treat a wide range of diseases, including various cancers. However, in drug development pipelines, the identification and subsequent optimization of effective small-molecule inhibitors can be a lengthy process. Traditional high-throughput screening methods are often laborious and cost-inefficient, with low hit rates [12]. In recent years, *in silico* methodologies including molecular docking and AI have emerged as indispensable cutting-edge tools that may have the potential to replace established HTS methods. Machine learning is remarkably efficient in identifying lead compounds within large databases, expediting the early stages of novel drug development. Deep Learning (DL) is a subcategory of machine learning based on artificial neural-networks. It has been shown to have applications across many fields of research, including various pharmaceutical sectors. In drug design, DL presents great promise because of its molecular predictive abilities and highly complex pattern recognition [13, 14, 15]. Deep Neural Networks (DNNs) have complex hierarchical structures, with several fully connected layers with many artificial neurons per layer. In computational drug discovery, a DNN receives input as molecular features. The network is trained using backpropagation, where errors from the output are propagated backward to adjust the weights of the neurons, thus optimizing predictive performance over many iterations. Incorporating DL models has the potential to increase the speed at which candidate compounds can be identified in a screening campaign.

**Figure 1.**
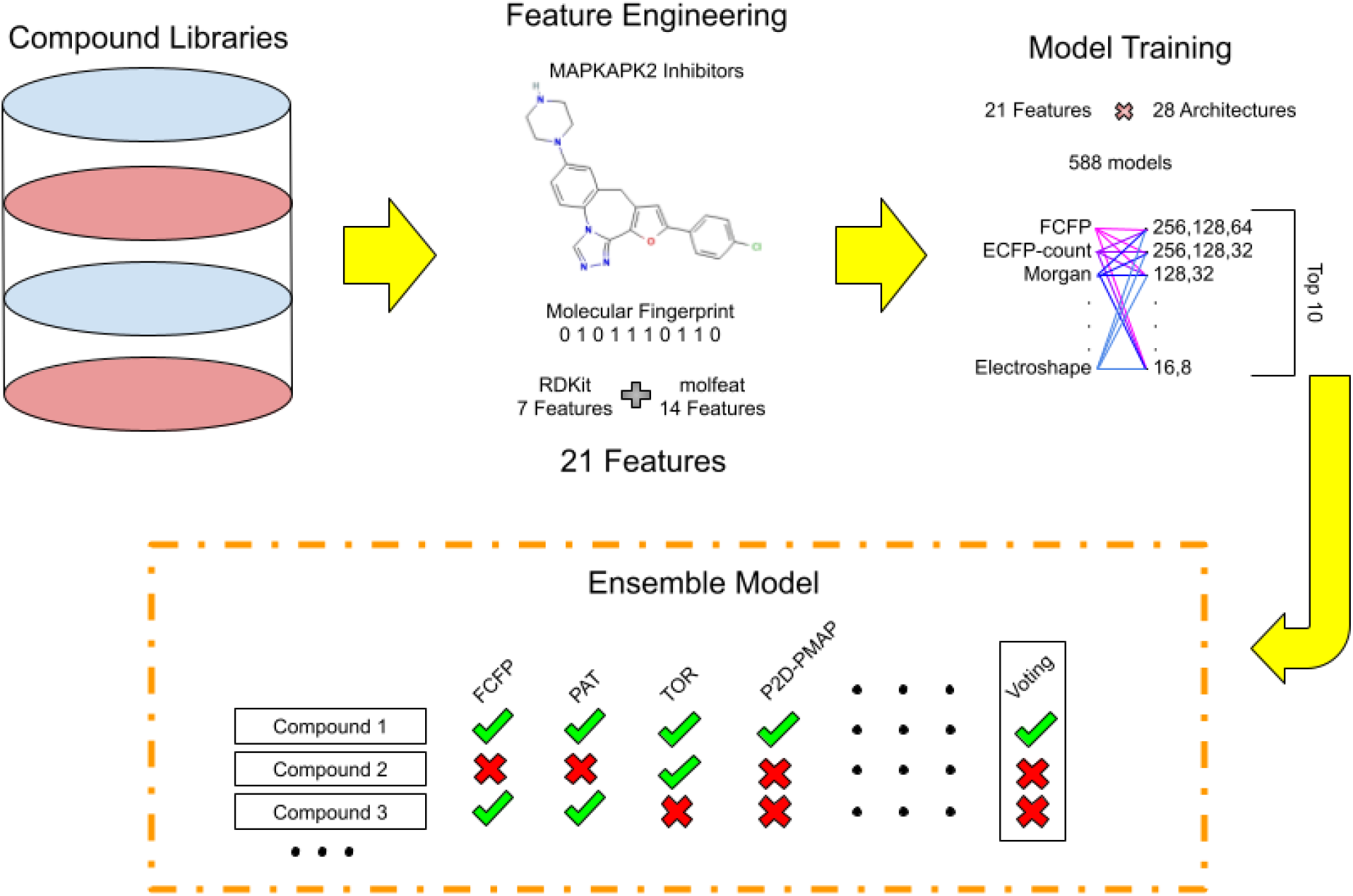
Workflow of the presented study.

## 3. Methods

### 3.1 Data Collection

A list of compounds with MAPKAPK2 bioactivity data was collected. Data points with bioactivity (MAPKAPK2 inhibition with IC50 values of under 10,000 nM) were labeled as “active” and data points with insufficient bioactivity (IC50 greater than 10,000 nM) were labeled as “inactive”. The dataset contained 2,950 compounds, with 840 classified as active and 2,110 as inactive. The dataset was then divided into training and testing sets in a 9:1 ratio, with the sets containing 2,655 and 295 compounds, respectively.

In addition, an Available Chemical Directory (ACD) dataset was obtained for model testing purposes. This compound set contained a total of 892 inactive compounds and was used to evaluate false positive rates.

### 3.2 Feature Engineering

Compounds were represented as molecular fingerprints and features using the RDKit and molfeat packages [16, 17]. The types of fingerprints selected from the RDKit package included Avalon, FeatMorgan, Layered, MACCS, Morgan (1024 bit), RDKit default, and Torsion. The types of fingerprints, featurizers and descriptors selected from the molfeat package included the 6 Potential Pharmacophore Points Chemically Advanced Template Search pharmacophore (CATS2D), Extended Connectivity Fingerprints-Count (ECFP-count), Electroshape, Extended Reduced Graph (ErG), Electrotopological state (Estate), Functional-class fingerprints (FCFP), Mordred, 2D Gobbi pharmacophore (pharm2D-gobbi), 2D Pmapper pharmacophore (pharm2D-pmapper), 3D Gobbi pharmacophore (pharm3D-gobbi), Pattern, ScaffoldKeys, SMILES extended connectivity fingerprint (SECFP), and Ultrafast Shape Recognition with Credo Atom Types (USRCAT). The number of molecular fingerprints, features, and descriptors totaled 21, with 9 running on RDKit and 12 running on molfeat.

For molecular featurizers that did not return binary bits from the SMILES string representations, the dataset was standardized (e.g. converted to z-scores) using StandardScaler on the Scikit-Learn API in Python before models were constructed using the dataset [18,19].

### 3.3 Model Construction

Multilayer Perceptron (MLP) models were built using the Keras API [20, 21]. Each MLP model used multiple layers of neurons between the input and output layers, with the weights of the neurons being updated by input data through backpropagation in accordance with the loss function. The binary cross-entropy loss function was used. Neuron layer sizes were varied between different models, with 28 different sets of layer sizes being tested for each of the 21 feature datasets. In each model, the activation function was ReLU for the input and hidden layers and Sigmoid for the output layer. The optimizer was Adam and default learning rate was used (0.001). The number of epochs was set to 100, and the batch size was kept at the default value (32). In addition, an early stopping strategy was implemented to prevent overfitting. The patience parameter was set to 10: when the loss function did not improve for 10 consecutive epochs, the model would cease training. A total of 588 models were trained using 21 different features, and each of the 588 models was evaluated via three strategies: 5-fold cross-validation, a testing dataset, and the ACD dataset consisting of solely inactive compounds. In 5-fold cross-validation, models were each evaluated on the training set (90% of the original dataset) 5 times, and the average performance metrics were taken. Then each model was retrained on the training set, and its predictive performance was evaluated against the testing dataset. Additionally, the models were also evaluated against the ACD dataset of inactive compounds, to test the false positive rate.

The performance of the models was evaluated by seven metrics: accuracy, precision, recall (sensitivity), specificity, F1 score, average precision score (AP), and area under the receiver operating characteristic curve (AUC-ROC). These metrics were calculated according to the following definitions:

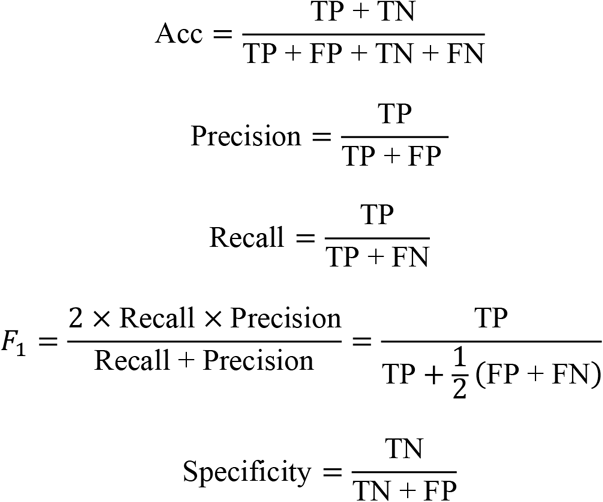

In the above formulas, The AP score and AUC-ROC were taken as the area under the Precision-Recall curve and the ROC curve, respectively.

The number of false positives identified in the inactive ACD dataset was used as an additional test for the validity of the models.

### 3.4 Model Selection and Voting

After a total of 588 models were created and evaluated, 10 of the top-performing models were selected. An ensemble model was constructed from the 10 models, and two voting systems were established. The first system returned the sum of the binary predictions of each model on a given set of molecules (0 for a negative prediction and 1 for a positive prediction). The second system returned the sum of the raw prediction probabilities of each model on the set of molecules. To prevent biased or imbalanced voting results, it was ensured that each of the selected models had been trained using datasets of distinct fingerprints/features. This ensured equal representation of featurizers in the ensemble model. The two voting systems were then evaluated using the test set of 295 compounds (0.1 of the original list of 2,950 compounds) and the inactive ACD compound list.

## 4. Results

### 4.1 Analysis and the Resulting Selection of Models

On the same feature type, 28 models with different DNN architectures were trained. The network layer sizes are shown below.

|  |  |  |  |
| --- | --- | --- | --- |
| 512,256 | 256,64 | 128,64 | 64,16 |
| 512,128 | 256,32 | 128,32 | 64,32,16 |
| 512,64 | 256,16 | 128,16 | 32,16 |
| 512,256,128 | 256,128,64 | 128,64,32 | 32,8 |
| 512,256,64 | 256,128,32 | 128,64,16 | 32,16,8 |
| 512,128,64 | 256,64,32 | 128,32,16 | 32,4 |
| 256,128 | 256,64,16 | 64,32 | 16,8 |

For each group of 28 models trained on the same feature type, metrics were compared between the models in that group, which only differed in network architecture (neuron layer sizes). The highest-performing model was determined by giving precedence to the accuracy metric, then considering other factors such as precision and AUC, to determine the validity of the model. One of the models, trained using the FCFP circular fingerprint, was found to have the highest overall metric scores (Table 1). This model had network layer sizes of 32 and 8. In this way, the model with the best-performing network architecture was determined from each group of models trained on the same feature type. The top ten determined models (Table 1) were chosen to form an ensemble model, which was then evaluated on the 0.1 test set and the inactive ACD compounds.

**Table 1.** Performance of selected models (5-fold cross validation). False Positive counts (rightmost column) on the inactive ACD dataset were recorded out of 892.

| Fingerprint/<br>Architecture | Accuracy | Precision | Recall | F1 | Specificity | AP | AUC | ACD<br>FP |
| --- | --- | --- | --- | --- | --- | --- | --- | --- |
| FCFP<br>[32,8] | 0.971 | 0.969 | 0.960 | 0.964 | 0.986 | 0.981 | 0.986 | 7 |
| ECFP-count<br>[256,32] | 0.971 | 0.967 | 0.960 | 0.963 | 0.984 | 0.981 | 0.986 | 8 |
| SECFP<br>[256,64] | 0.966 | 0.964 | 0.953 | 0.958 | 0.984 | 0.974 | 0.979 | 10 |
| Pharm2D-Gobbi<br>[64,32] | 0.965 | 0.962 | 0.952 | 0.957 | 0.983 | 0.972 | 0.980 | 8 |
| FeatMorgan<br>[256,128] | 0.962 | 0.954 | 0.953 | 0.953 | 0.975 | 0.973 | 0.981 | 6 |
| Pharm2D-Pmapper<br>[128,64] | 0.962 | 0.957 | 0.950 | 0.953 | 0.978 | 0.971 | 0.980 | 2 |
| Morgan<br>[128,64] | 0.961 | 0.957 | 0.946 | 0.952 | 0.981 | 0.974 | 0.981 | 3 |
| Pattern<br>[512,256,<br>128] | 0.959 | 0.955 | 0.944 | 0.949 | 0.979 | 0.970 | 0.982 | 0 |
| Torsion<br>[512,128,64] | 0.959 | 0.948 | 0.953 | 0.950 | 0.967 | 0.971 | 0.981 | 4 |
| Layered<br>[512,128] | 0.957 | 0.947 | 0.947 | 0.947 | 0.970 | 0.971 | 0.982 | 1 |

**Figure 2.**
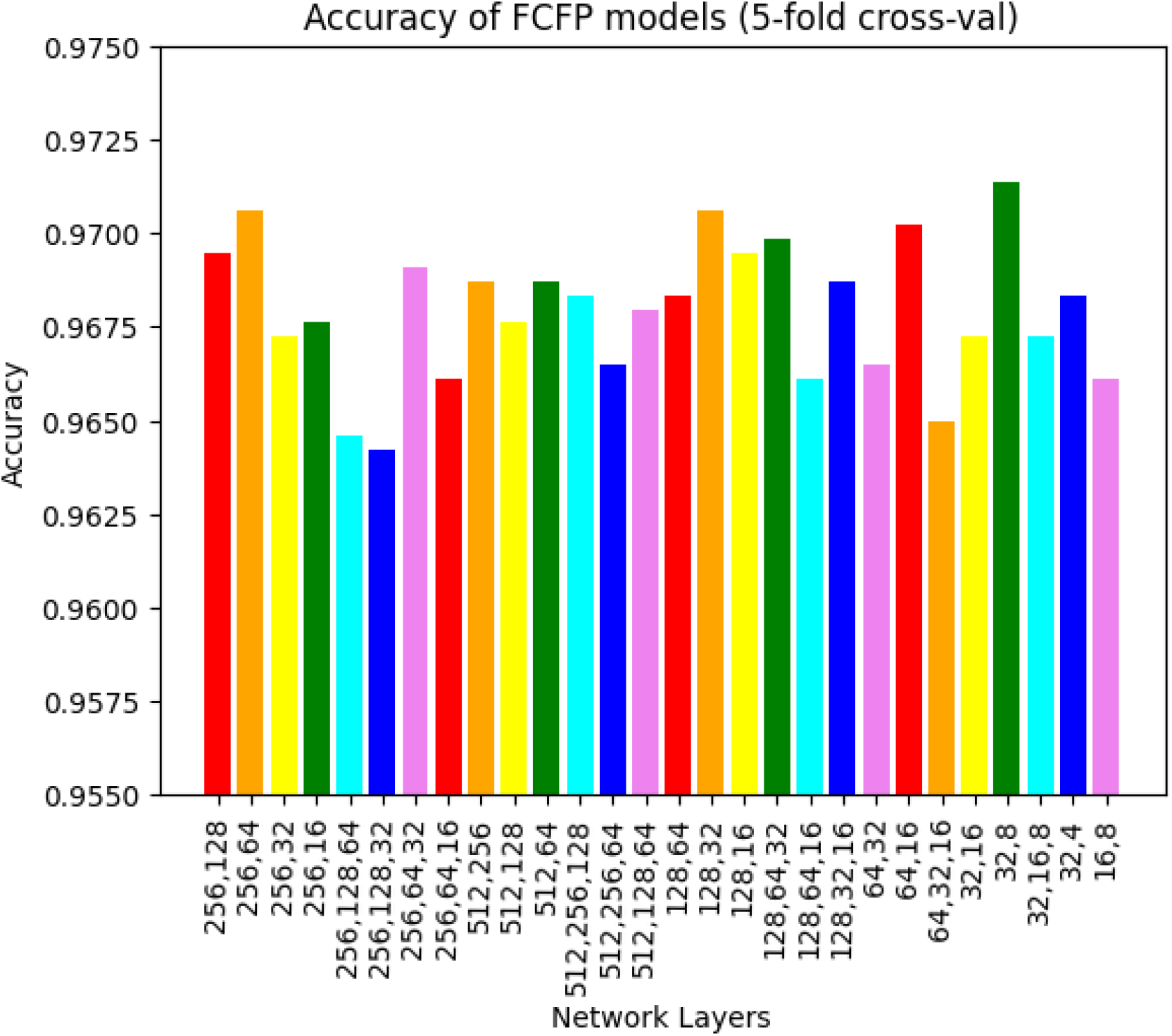
Comparison of accuracy between models trained on the same feature. FCFP models were used as an example here. The 32,8 network architecture was thus chosen as the highest-performing in this group.

**Figure 3.**
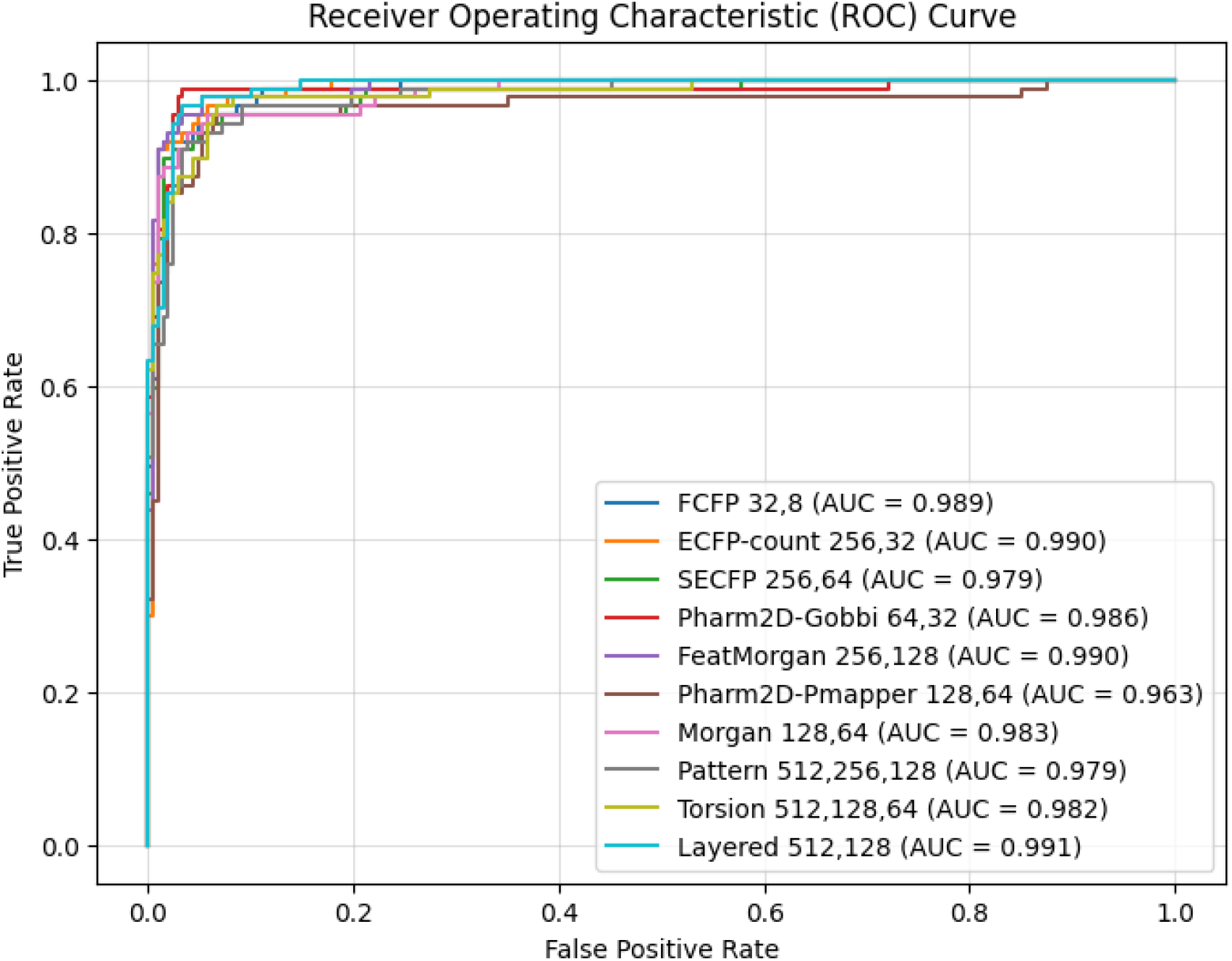
Comparison of Receiver Operating Characteristic Curves on Test Set. High AUC-ROC values of over 0.96 indicated favorable performance of the selected models for MAPKAPK2 inhibitor classification.

### 4.2 Evaluation of Ensemble Models

Ensemble models were evaluated on the test set, which contained 295 compounds. Different consensus “voting” methods were examined. Per query molecule in the test set, a set number of the 10 separate selected models was required to identify the molecule as having MAPKAPK2 bioactivity. This number ranged from 5 to 10 (Table 2). This was the first voting system, which will here be denoted “binary voting”. It was discovered that the best-performing ensemble model required 7 out of 10 possible votes from the constituent selected models. It was observed that as the strictness of the voting criteria increased (requiring more models to select a given molecule), the accuracies and precisions of the ensemble models initially increased, then later decreased after peaking at 0.969 and 0.964, respectively, when the number of votes required was 7 (Table 2). Moreover, the false positive rate ceased to decrease at a voting threshold of 7 (remaining constant at 0.014), while the Recall metric sharply declined from 0.931 to 0.793 as the voting threshold was increased to 10. This demonstrated that when the voting threshold was too high, or too strict, the ensemble model could not reliably identify active MAPKAPK2 inhibitor molecules. In this way, the voting threshold that gave the ensemble model the best performance was determined to be 7 out of the 10 possible votes from the 10 selected individual models. A reasonable range of thresholds was determined to be approximately between 6 and 8.

**Table 2.**
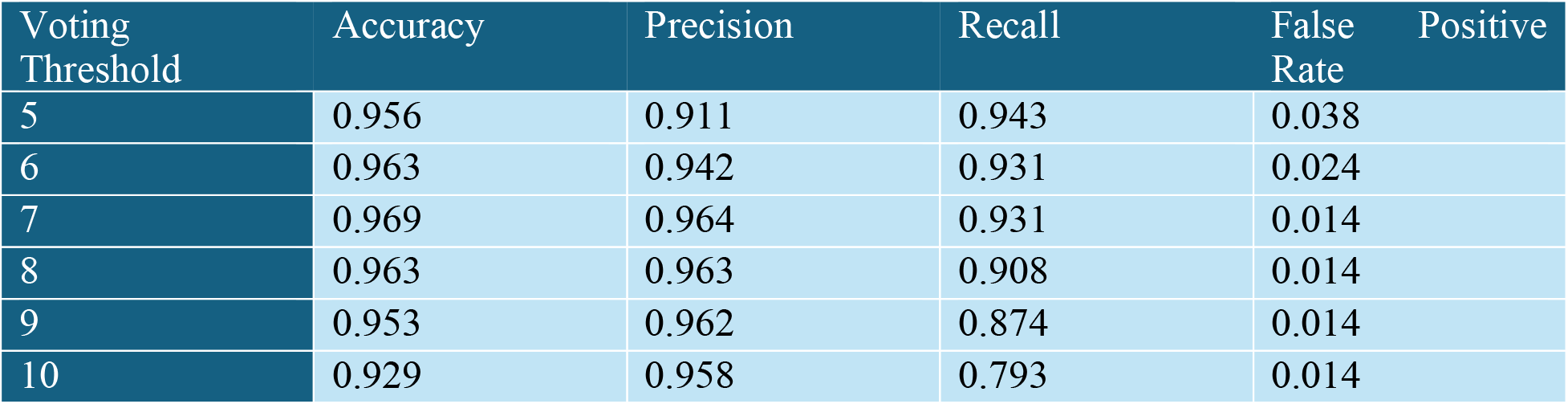
Evaluation of different voting thresholds in ensemble model. The optimal voting requirement for such a consensus “voting” system was thus determined to be 7 out of 10 possible votes.

A second voting system was explored, which will be denoted “probability sum”. Thresholds were set as the sum of models’ raw prediction probabilities on each molecule in the dataset. The thresholds would ideally be tested in a non-discrete manner, as predictive probabilities may take on infinitely many values between 0 and 1 inclusive. However, as evaluations were done on a finite test set with only 295 compounds, varying the probability sum threshold by some infinitely small amount ε would be unlikely to change the classification of any given molecule by the ensemble model. It was deduced that minimal adjustments in the strictness of the voting system would produce negligible changes in ensemble model performance for this specific test set. Thus, probability sum thresholds were chosen in increments of 0.5, within a reasonable range of 5.5 to 8 (Table 3). Ensemble model performance across different probability sum thresholds showed similar trends as performance across different binary voting thresholds (Table 2). In this case, the ensemble model showed the best results (had the best scores across all examined metrics) when the probability sum threshold was set to 6.5. These performance results were, in fact, identical to the results produced when the binary voting threshold was set to 7 (Table 2).

**Table 3.** Evaluation of different probability sum thresholds in ensemble model. The optimal threshold for raw probability sums was determined to be 6.5 for the test set.

| Probability Sum Threshold | Accuracy | Precision | Recall | False Positive Rate |
| --- | --- | --- | --- | --- |
| 5.5 | 0.966 | 0.953 | 0.931 | 0.019 |
| 6 | 0.966 | 0.953 | 0.931 | 0.019 |
| 6.5 | 0.969 | 0.964 | 0.931 | 0.014 |
| 7 | 0.959 | 0.963 | 0.897 | 0.014 |
| 7.5 | 0.953 | 0.962 | 0.874 | 0.014 |
| 8 | 0.946 | 0.961 | 0.851 | 0.014 |

The ensemble model was then tested against the inactive ACD dataset, and the results compared with individual models. Notably, when predictions for all 892 of the inactive molecules were made, the ensemble model only returned predictive binary voting scores of 3 or greater for 4 of the 892 molecules, and there were no scores over 5. Using the probability sum voting method, the ensemble model returned scores of 3 or greater for 8 of the 892, and the maximum probability sum score was about 5.093 (truncated to three decimal places). These prediction results indicated that if the binary voting threshold and probability sum threshold required to indicate MAPKAPK2 bioactivity were fixed at 7 and 6.5, respectively, no false positives would be identified. Thus, the ensemble model has high precision and could be used to successfully curate a MAPKAPK2-inhibition-focused compound library.

**Figure 4.**
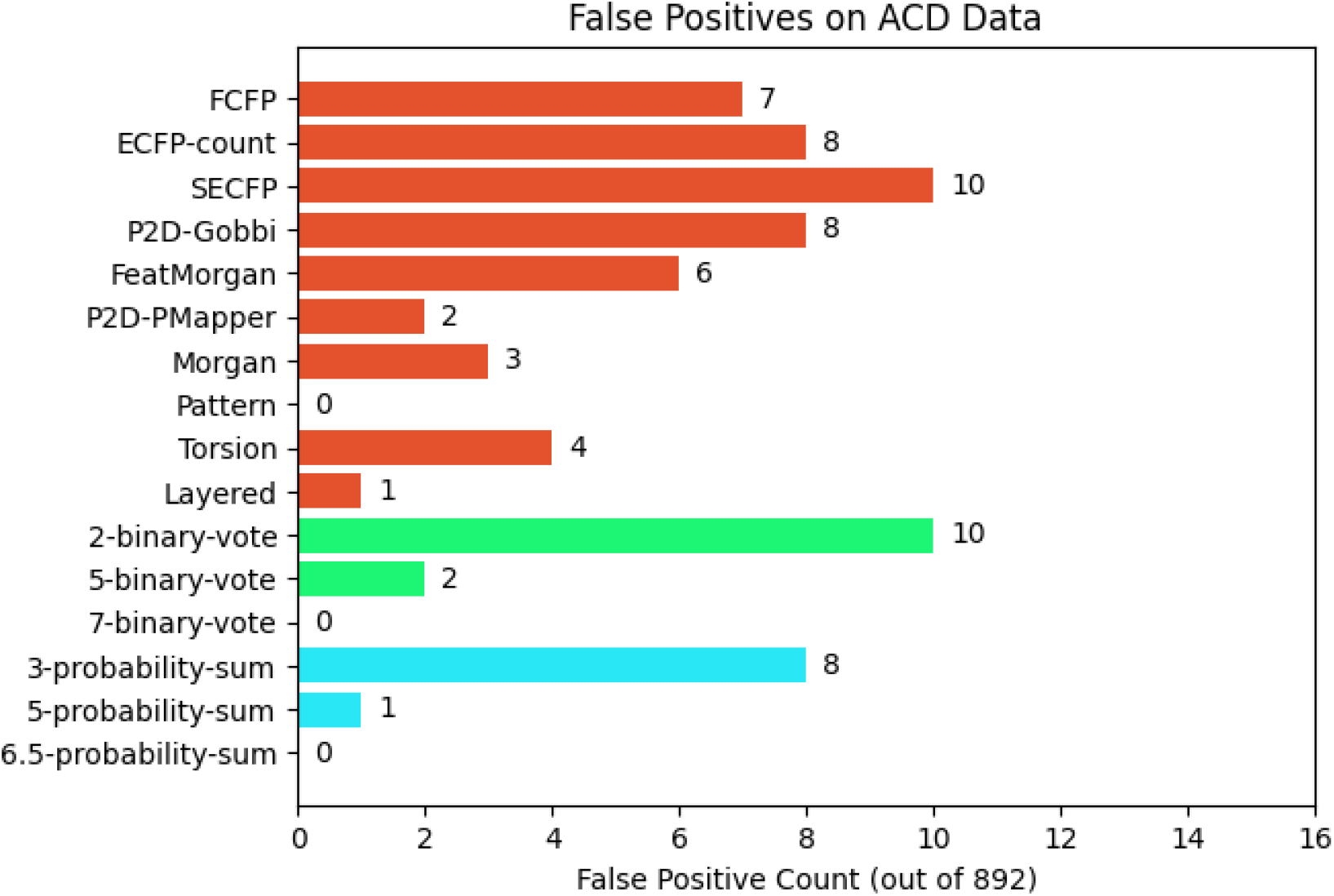
Comparison of False Positives identified in ACD dataset between individual and ensemble models. It was observed that either requiring 7 binary votes or a total probability sum of 6.5 produced no false positives on this compound list, a satisfactory result given that 9 of the 10 individual models gave false positive predictions on the ACD set.

## 5. Discussion

MAPKAPK2 is an important protein kinase in the p38 MAPK pathway that is relevant in cancer, arthritis, and many inflammatory diseases. Inhibition of this protein kinase has been shown to have benefits in treating cancers among other diseases. In this study, a ligand-based ensemble model incorporating ten different molecular features was successfully created and optimized. Voting systems were also examined, and two of the best-performing voting methods were established. The preparation of a screening library is a crucial step for successful screening campaigns. The ensemble model and voting systems can next be used to screen compound sets for MAPKAPK2 bioactivity to create a MAPKAPK2-focused library for further screening analysis and hit compound identification using techniques such as molecular docking. Additionally, feature importance analysis can also reveal the crucial features in the individual models that comprise the established ensemble model. Identified novel hit compounds will function as additional structures of study for the development of new MAPKAPK2 therapeutics. Given available bioactivity training data, ensemble models can be created for other protein kinase targets in addition to MAPKAPK2, highlighting a viable strategy for the identification of compounds with bioactivity within large databases as an initial step in the drug development pipeline. In conclusion, this study revealed an effective method for the integration of many molecular features and descriptors in creating ensemble models for the prediction of novel protein kinase inhibitors.

